# Zinc Determines Dynamical Properties and Aggregation Kinetics of Human Insulin

**DOI:** 10.1101/2020.07.02.184903

**Authors:** K. Pounot, G. W. Grime, A. Longo, M. Zamponi, D. Noferini, V. Cristiglio, T. Seydel, E. F. Garman, M. Weik, V. Foderà, G. Schirò

## Abstract

Protein aggregation is a widespread process leading to deleterious consequences in the organism, with amyloid aggregates being important not only in biology but also for drug design and biomaterial production. Insulin is a protein largely used in diabetes treatment and its amyloid aggregation is at the basis of the so-called insulin-derived amyloidosis. Here we uncover the major role of zinc in both insulin dynamics and aggregation kinetics at low pH, where the formation of different amyloid superstructures (fibrils and spherulites) can be thermally induced. Amyloid aggregation is accompanied by zinc release and the suppression of water-sustained insulin dynamics, as shown by particle-induced X-ray emission and X-ray absorption spectroscopy and by neutron spectroscopy, respectively. Our study shows that zinc binding stabilizes the native form of insulin by facilitating hydration of this hydrophobic protein and suggests that introducing new binding sites for zinc can improve insulin stability and tune its aggregation propensity.

**Statement of Significance:** Localized amyloidosis occurs at insulin injection sites for diabetes treatment, leading to deleterious inflammations known as insulin-derived amyloidosis. Amyloid superstructures are also promising candidates in the field of biomaterials. Here we revealed that zinc, coordinated to insulin in the native form, is released upon amyloid aggregation, when insulin forms superstructures known as fibrils and spherulites. Zinc release leads to a full suppression of functionally essential protein dynamics through a modification of the protein’s hydration properties and completely modifies insulin amyloid kinetics. The results suggest that changes in protein hydration upon zinc binding/release modifies both stability and dynamics of insulin and might then be a general strategy to control protein stability and tune protein aggregation into amorphous and ordered superstructures.

## Introduction

*In vitro* under destabilizing conditions, a large number of proteins aggregate and form specific ordered superstructures (1). These can be amyloid fibrils, characterized by a typical cross-beta pattern (2, 3) or larger, micrometric assemblies of amyloid nature, like spherulites (4, 5). The variety of amyloid species poses a challenge for pharmaceutical drug development, where protein-based products have to be optimized while controlling and characterizing any by-product particles. Indeed, due to the simultaneous occurrence of different species, identifying and isolating each individual species is a highly demanding task, still remaining a *conditio sine qua non* for the quality control of the final product (6). On the other hand, having access to a number of markedly different protein self-assembled species offers a unique opportunity for the development of a rational platform for the design of bio-inspired materials for applications ranging from bio-sensing and tissue engineering to drug delivery (7). Finally yet importantly, a large-scale morphological variability of aggregates is also observed in the context of amyloid-related pathologies. This variability potentially determines different etiological subtypes of diseases (8–10), making it even more challenging to univocally establish a connection between aggregate structure/morphology and the onset/progression of the diseases. In all the above fields, it is then crucial to disentangle the concurring aggregation pathways and isolate the different species. Indeed, identifying the different aggregates would allow the mechanisms of formation of the different species to be mapped, as well as unraveling how the structure-function-dynamics relationship is perturbed by amyloid formation for each of the different aggregation states.

Insulin, a 5.7 kDa hormone consisting of two mainly alpha-helical chains (A and B) linked by disulfide bonds, can form amyloid fibrils (11, 12) and spherulites (13). The two types of aggregate coexist at low pH when human insulin samples are thermally destabilized (14, 15). Subcutaneous insulin injections are largely used during insulin therapy for diabetic patients. Near injection sites, localized amyloidosis has been found that can lead to deleterious repeated inflammations, known as the insulin-derived amyloidosis (IDA) (16–19) or also “insulin ball” (20) or amyloidoma (21). The number of IDA reported cases has increased significantly in the last years (22). When IDA occurs, it leads to poor glycemic control and also to catheter occlusion in the case of continuous infusion. Recent *in vitro* studies on the stability of insulin formulations used to treat type 1 diabetes have shown that amyloid formation occurs either at neutral pH or below the insulin isoelectric point (pH < 5.2) (23). An alternative way recently explored for insulin delivery to diabetic patients is the administration by the oral route, that could potentially overcome the drawbacks of insulin injection reducing the number of injections needed and the risk of side effects (24). However, when administered orally, insulin first arrives at the stomach, where pH is between 1.2 and 3.0, which is the range where, *in vitro*, insulin forms both amyloid fibrils (11) and spherulites (13). It is also known that diabetes is commonly associated with neurodegenerative diseases (25) and insulin signaling impairment - for which insulin aggregation might play a role - was shown to promote neurodegeneration (26, 27). Furthermore, insulin stability is a major concern in the pharmaceutical industry for production and storage (8–10). As a consequence, understanding the key factors involved in insulin self-assembly is highly desired for the optimization of downstream product processing, as well as for a more exhaustive mapping of the multiplicity of different structures occurring in its aggregation reaction. The insulin aggregation pathway starts from a mixture of oligomeric forms. The oligomerization state depends on the solution conditions, with mainly a dimer-monomer mixture occurring at low pH (28) and a hexamer species at higher pH. However, insulin amyloid species can be formed independent of the details of the early oligomerization process (29) and such aggregates are Thioflavin T (ThT)-positive, indicating the presence of a cross-beta fibrillar structure as is found for toxic amyloid proteins such as tau or α-synuclein (2, 3). Several factors are reported to affect the aggregation process of insulin. Approaches involving sedimentation kinetics measurements (30), optical microscopy imaging (14), and small angle X-ray scattering (31) have revealed the role of both insulin and salt concentration and of pH in the aggregation process. In the context of amyloid diseases, the pathological impact of metal ions in the alteration of amyloidogenesis is now accepted (32). In the specific case of insulin, zinc ions (Zn^2+^) are essential components for protein expression and activity (33). It has been shown that Zn^2+^ slows down fibrillation of monomeric insulin at physiological pH: this result has been used to propose that Zn^2+^ co-secreted from pancreatic β-cells protects the organism from the formation of non-native insulin oligomers and aggregates (34). However, Zn^2+^ has been found to be coordinated in both functional and aggregated bovine insulin at physiological pH (35).

More generally, the significant influence of metal ions on aggregation kinetics, pathway, and aggregate morphology has been reported for several amyloid systems (36–38). As a matter of example, zinc and copper ions increase the stability of Aβ oligomers and reduce the stability of Aβ fibrils, the effect on fibrils being specific to the type of ion (39). Ions can also dramatically affect the balance between the numbers of native multimers and monomers (40), which are prone to form amyloid-like fibrils in systems containing GAPR-1 and heparin (41). Notwithstanding pronounced metal ion-related effects are clearly detected in other amyloid-forming systems (37, 42–46), the mechanism at play remains elusive. Metal coordination at the level of single protein molecules certainly changes the overall charge, protein structure and/or hydration, eventually influencing the dynamics of the single protein molecule (47, 48). How these changes modify the physico-chemical properties of protein aggregates, however, remains unclear.

Recently, several pieces of evidence have been reported on the relationship between amyloid aggregation and the dynamical properties of aggregating proteins. Measurements of intramolecular reconfiguration dynamics in different pathological amyloid proteins suggested a direct correlation between internal dynamics and aggregation propensity and kinetics (49, 50). Neutron scattering results obtained on different proteins showed that upon amyloid aggregation either the protein (51) or its hydration water (52) can show an increased mobility at temperatures above the so-called dynamical transition (53).

Here, we used neutron scattering to characterize the dynamics of human insulin in its native state and in both its fibril and spherulite amyloid forms. In order to characterize the internal dynamics of both fibrils and spherulites, we first screened and selected the conditions for isolating samples containing either predominantly fibrils or predominantly spherulites. Neutron scattering revealed a complete suppression of functionally relevant internal protein dynamics in both insulin fibrils and spherulites. An analysis based on microbeam Particle Induced X-ray Emission (μPIXE) and X-ray absorption spectroscopy (XAS), combined with a protein conformational characterization, unexpectedly revealed that the protein rigidification is due to the release, upon amyloid aggregation, of the Zn^2+^ coordinated to insulin in the native state (33). A crucial role of Zn^2+^ in determining insulin aggregation kinetics was then unveiled by fluorescence spectroscopy.

## Materials and Methods

### Sample preparation: native and EDTA treated insulin

Crystalline powder of human insulin was obtained from Novo Nordisk and stored at −20°C before use. In order to dissolve crystals to obtain monomeric samples, the powder was dissolved by adding 5 μl of HCl to a suspension in H_2_O until the solution was clear. After flash-cooling in liquid nitrogen and freeze-drying, electron microscopy and X-ray powder diffraction experiments were used to check for the presence of any crystals – protein or salt – and for aggregates. For the washed and EDTA treated samples, the freeze-dried solutions were dissolved with and without EDTA, respectively. The pH was then set to the insulin isoelectric point (pI = 5.2) to perform washing steps using several cycles of centrifugation at 18000 rpm for 20 minutes and resuspension in pure D_2_O. Finally, flash-cooling and freeze-drying were performed prior to D_2_O hydration for the neutron experiments.

### Sample preparation: insulin fibrils and spherulites

The conditions for preparing pure fibril and pure spherulite samples are described in the main text, as they were obtained and optimized in the present work. Prior to neutron experiments, fibrils and spherulites were washed using several cycles of centrifugation at 18000 rpm for 20 minutes and resuspension in pure D_2_O to remove any salt and contaminants. They were then flash-cooled and freeze-dried as for the monomeric samples.

### Incoherent elastic neutron scattering

All data were recorded on the SPHERES backscattering spectrometer (proposal P14013, 14-20^th^ June, 2018) (54, 55) operated by JCNS at the Heinz Maier-Leibnitz Zentrum (MLZ) in Garching, Germany. Freeze-dried insulin powders were incubated with P_2_O_5_ for 24 hours in a desiccator to further remove water. In the case of lysozyme, such a procedure resulted in a residual presence of four waters per protein molecule. The insulin powders were then weighed and re-hydrated by vapor-exchange with pure D_2_O to reach a hydration level h = 0.44 D_2_O/protein w/w. Hydrated protein powders were then sealed in aluminum flat neutron cells (sample thickness=0.3 mm) using 1 mm indium wire from Alfa Aesar. Samples were weighed before and after the neutron experiment to verify that they were correctly sealed and no water was lost. The cell was mounted in a cryostat at room temperature, and the temperature was subsequently lowered to 10 K within approximately 2 hrs. The elastic scattering signal was recorded at an energy resolution of 0.66 μeV (FWHM), employing Si(111) monochromator and analyzer crystals in exact backscattering geometry (for large scattering angles), corresponding to an incident wavelength of λ = 6.27 Å. The elastic signal was recorded while continuously increasing the sample temperature from 10 to 300 K at a rate of 0.2 K/min and binned together to 0.05 K steps. Raw data were pre-processed - i.e. normalized to the detector efficiency and to the incident beam intensity recorded by a so-called monitor device - using the SLAW software available on facility computers (http://apps.jcns.fz-juelich.de/slaw). With the elastic signal arising mainly from the incoherent scattering of the hydrogen atoms, the mean square displacements (MSD) of hydrogen atoms can be obtained using the well established Gaussian approximation (56, 57). We performed this analysis using custom-made python scripts (http://github.com/kpounot/nPDyn), with the Gaussian approximation verified for the momentum transfer (q = 4πλ^-1^sin(θ/2), with θ being the scattering angle) range 0.6-1.2 Å^-1^. Using higher terms in cumulant expansion leading to a corrected Gaussian model (57) or using a gamma distribution based model (58) gave similar results but with higher numerical instability during the fitting procedure.

### Neutron diffraction

Data was recorded on the D16 instrument at the ILL in Grenoble, France. The same samples within the sealed aluminum cells as used for measuring the incoherent elastic neutron scattering were mounted in an Orange cryostat to measure the data at two temperatures, 300 K and 200 K. The diffracted beam was measured over an angular range 12 - 112.5° which corresponds to a *q*-range 0.05 - 2.5 Å^-1^. Neutron data were corrected for the empty cell scattering, the ambient room background and the nonuniform detector response. The transmission and the thickness of the sample were also taken into account. The 2D scattering intensities were normalized in absolute units with a standard calibration and radially integrated to obtain 1D diffraction patterns.

### Microbeam Particle Induced X-ray Emission (μPIXE)

Zinc stoichiometries in the different insulin samples were measured using μPIXE analysis in combination with simultaneous Rutherford backscattering analysis (RBS) to allow correction for sample matrix effects (59, 60). For μPIXE measurements of proteins, sulfur acts as an internal standard. The measurements were carried out at the Ion Beam Centre, University of Surrey, UK (61). A 2.5 MeV proton beam of 1.5 μm in diameter was used to induce characteristic X-ray emission from dried insulin sample droplets (volume per droplet ~0.1 μl) under vacuum. The X-rays were detected using a solid state lithium drifted silicon detector and backscattered protons were detected using a silicon particle detector. By scanning the proton beam in *x* and *y* over the dried sample, spatial maps were obtained of all elements heavier than magnesium present in the sample. Quantitative information was obtained by collecting 3 or 4 point spectra from each droplet. PIXE spectra were analyzed with GUPIX (62) using the matrix composition derived from the simultaneous RBS spectrum. Data processing is carried out using the data acquisition software OMDAQ-3 (Oxford Microbeams Ltd, UK) to extract the relative amount of each element in the sample.

### X-ray absorption spectroscopy

#### Data collection

X-ray absorption spectra were collected at the zinc K-edge (9660.75 eV) on the EXAFS station (BM26A) of the Dutch-Belgian beamline (DUBBLE) (63) at the European Synchrotron Radiation Facility (ESRF) in Grenoble, France. The energy of the X-ray beam was tuned by a double-crystal monochromator operating in fixed-exit mode using a Si(111) crystal pair. Three different samples were put in separate glass capillaries and measured in fluorescence mode at ambient temperature and pressure: i) a zinc solution prepared by dissolving 4 mg of ZnSO_4_ in 10 ml of 0.25 M NaCl solution at pH 1.8, ii) a native insulin solution prepared by dissolving 10 mg of lyophilized insulin powder (Novo Nordisk) in 1 ml of 0.25 M NaCl solution at pH 1.8, and iii) the same solution as in ii) incubated at 60° C for about 24 h to induce amyloid aggregation. The EXAFS spectra, three scans per sample with a new solution for each scan, were energy-calibrated, averaged and further analyzed using GNXAS (64, 65).

#### Data analysis

In the GNXAS approach, the local atomic arrangement around the absorbing atom is decomposed into model atomic configurations containing 2,…, n atoms. The theoretical EXAFS signal **χ**(k) is given by the sum of the n-body contributions **γ**^2^, **γ**^3^,…, **γ**^n^, which take into account all the possible single and multiple scattering (MS) paths between the n atoms. The fitting of **χ**(k) to the experimental EXAFS signal allows refinement of the relevant structural parameters of the different coordination shells; the suitability of the model is also evaluated by comparison of the experimental EXAFS signal Fourier transform (FT) with the FT of the calculated **χ**(k) function. The coordination numbers and the global fit parameters that were allowed to vary during the fitting procedure were the distance R(Å), Debye-Waller factor (**σ**^2^) and the angles of the **γ**^n^ contributions which were defined according to atomic structural models constructed using the VMD software(66). Zinc was solvated based on a published refined structure (67). For the model containing histidine, the residue was positioned by replacing one coordinating water molecule with it. Weak distance constraints were used, so that water and histidine could compete to coordinate the zinc atom. Simulations were then performed using NAMD 2.13 (68) with the TIP3P model for water (69) and the CHARMM36 force field (70). The Nose-Hoover-Langevin piston algorithm maintained a constant pressure (71, 72), and the stochastic velocity rescaling algorithm controlled the temperature (73). Bonds with hydrogen atoms were constrained using the SHAKE algorithm (74) with a force constant k set to 2, and the Verlet-I/r-RESPA multiple-time step scheme (74–76) integrated the equations of motion, with time steps of 2 fs for long-range nonbonded forces, and 1 fs for short-range and bonded forces. Electrostatic interactions were computed using the smooth Particle Mesh-Ewald (PME) sum (77), with a cutoff set to 10 Å, a switching function starting at 8 Å, and a pairlist distance of 14 Å was used. A snapshot of the simulation run was then used to extract coordinates that were processed by GNXAS. According to the GNXAS approach, we used a single two body configuration **γ**^2^ for the Zn-L (L=O or N) distance. In order to fit the higher shell contributions two **η**^3^ corresponding to the Zn-O-C and Zn-O-N three-body configurations, were used (see inset in Figure 5c). Importantly, due to the high value of the vertex angle in the Zn-O-C and Zn-O-N configurations, which are connected to the aromatic ring present in the histidine, the signals of theses shells are enhanced because of the multiple scattering effect, which is considered in the analysis. For these shells, whose Zn-O distance is the first shell, the only free parameter needed was the vertex angle **θ**.

**Figure 1.**
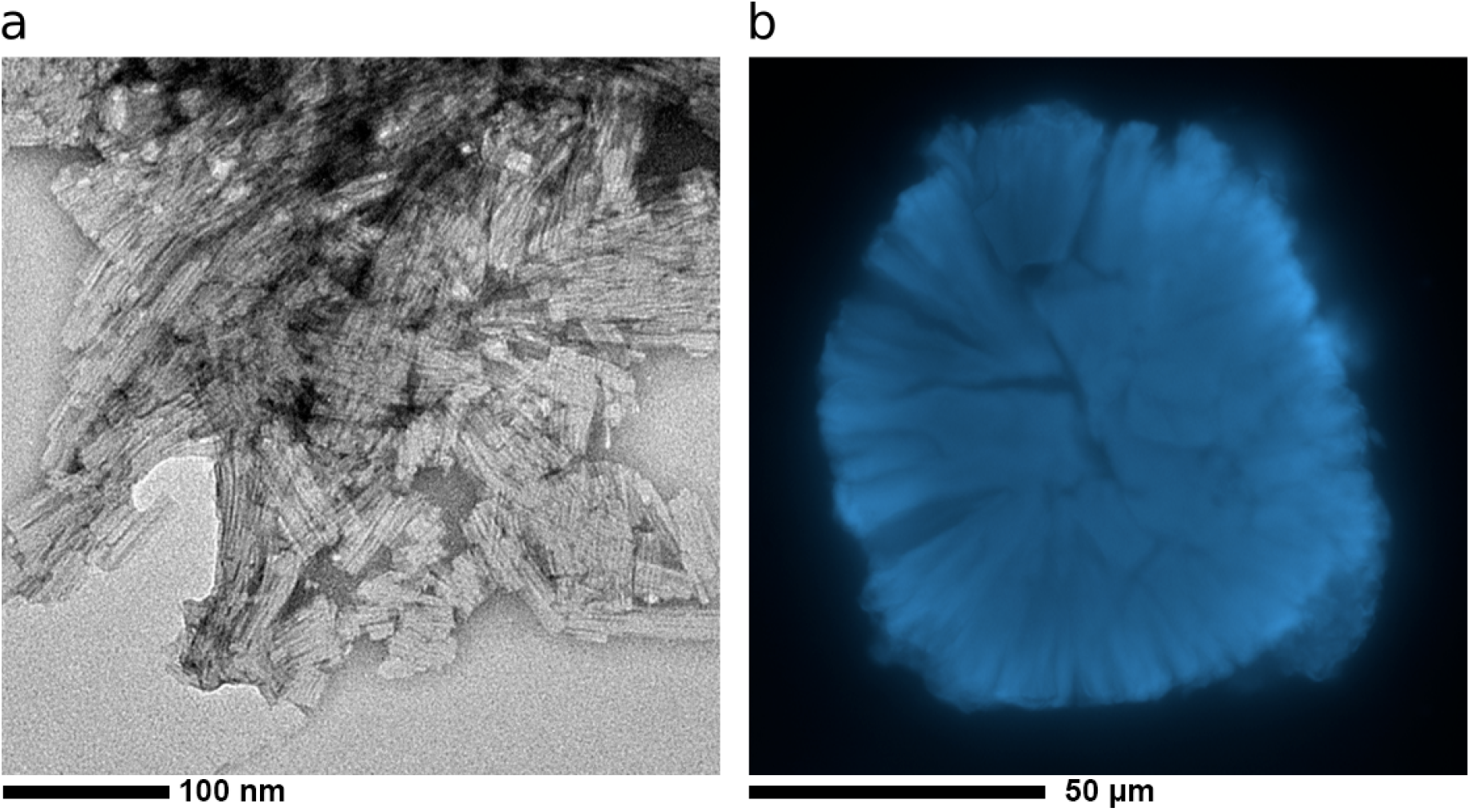
Morphology of insulin amyloid aggregates. **a** Electron microscopy of freeze-dried insulin fibrils. **b** Confocal microscopy of a freeze-dried insulin spherulite.

**Figure 2.**
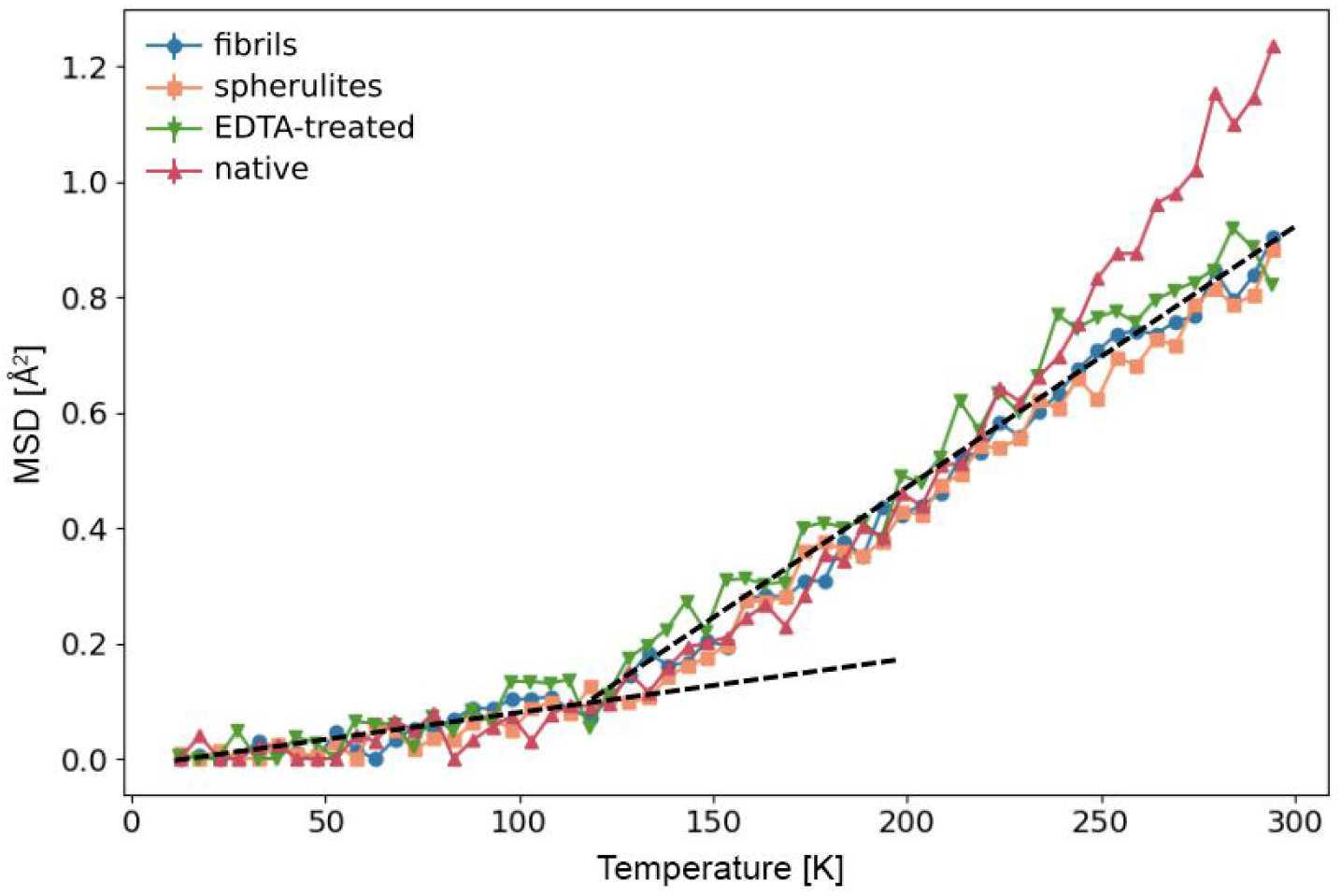
Amyloid aggregation suppresses the protein dynamical transition of hydrated insulin. Mean square displacements (MSD) measured as a function of temperature by elastic incoherent neutron scattering on the backscattering spectrometer SPHERES (MLZ, Garching) for native insulin (red triangles), amyloid fibrils (blue circles) and spherulites (orange squares) and for insulin treated with EDTA (green triangles).

**Figure 3.**
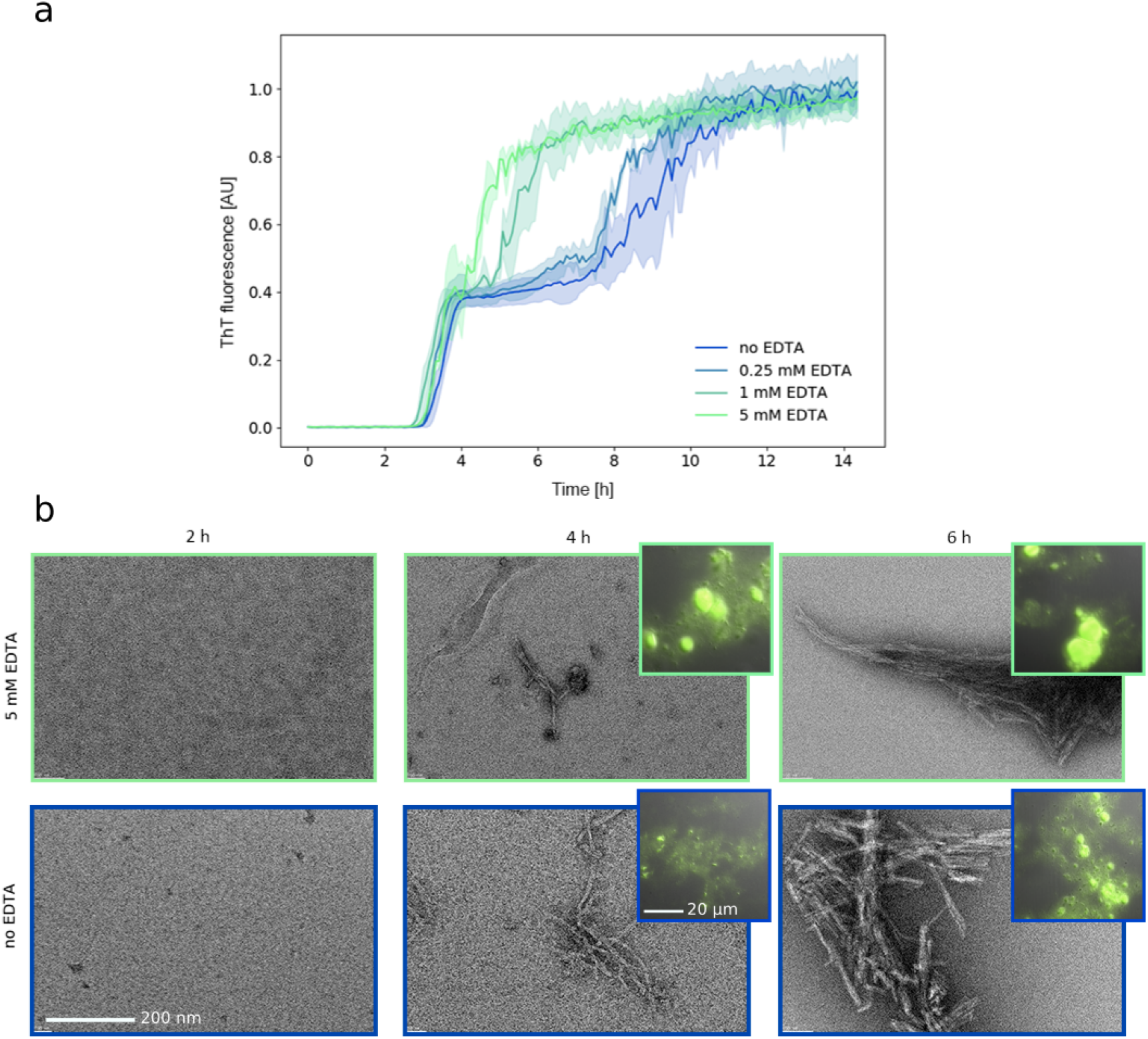
EDTA modifies insulin aggregation kinetics. **a** Insulin aggregation kinetics at 60° C as probed by Thioflavin T fluorescence as a function of EDTA concentration. The color shaded areas indicate the standard deviation for three independent aggregation processes. **b** Transmission electron microscopy (fibrils) and fluorescence microscopy (spherulites, insets) images acquired during the kinetics without EDTA (blue curve in **a**, blue contoured images in **b**) and at 5 mM EDTA (light green in **a**, light green contoured images in **b**). Micrographs were acquired after 2 h (left), 4 h (middle) and 6 h (right) of incubation at 60° C and showed that fibrils and spherulites were formed roughly at the same time, with more numerous and larger spherulites appearing in the presence of EDTA.

**Figure 4.**
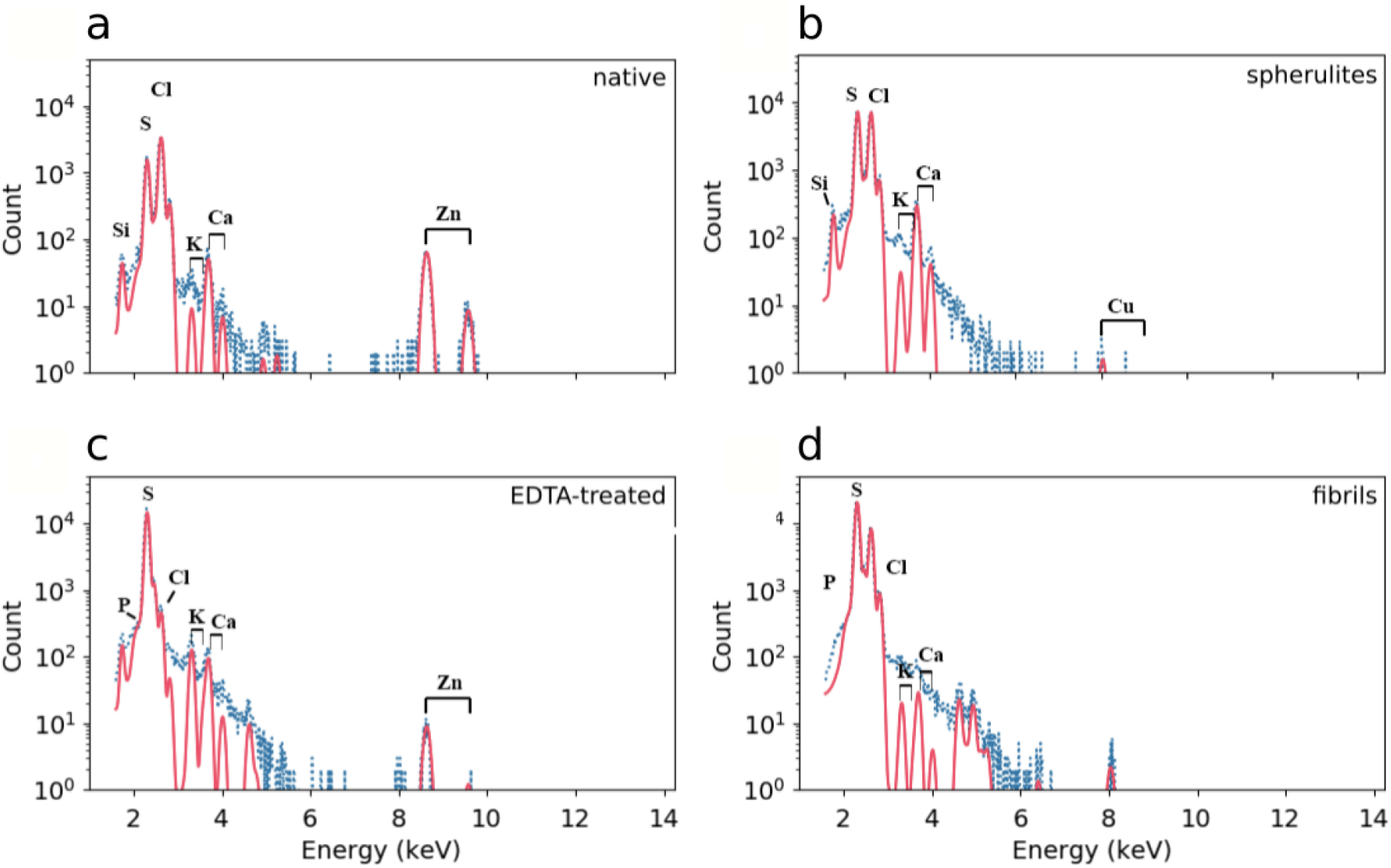
Insulin loses coordinated zinc upon amyloid aggregation. Proton Induced X-ray Emission (PIXE) spectra (blue points) and fit to the characteristic peaks (red lines) for (**a**) native insulin, (**b**) insulin spherulites, (**c**) EDTA treated insulin, and (**d**) insulin fibrils. The theoretical positions for X-ray emission maxima are indicated in all cases.

**Figure 5.**
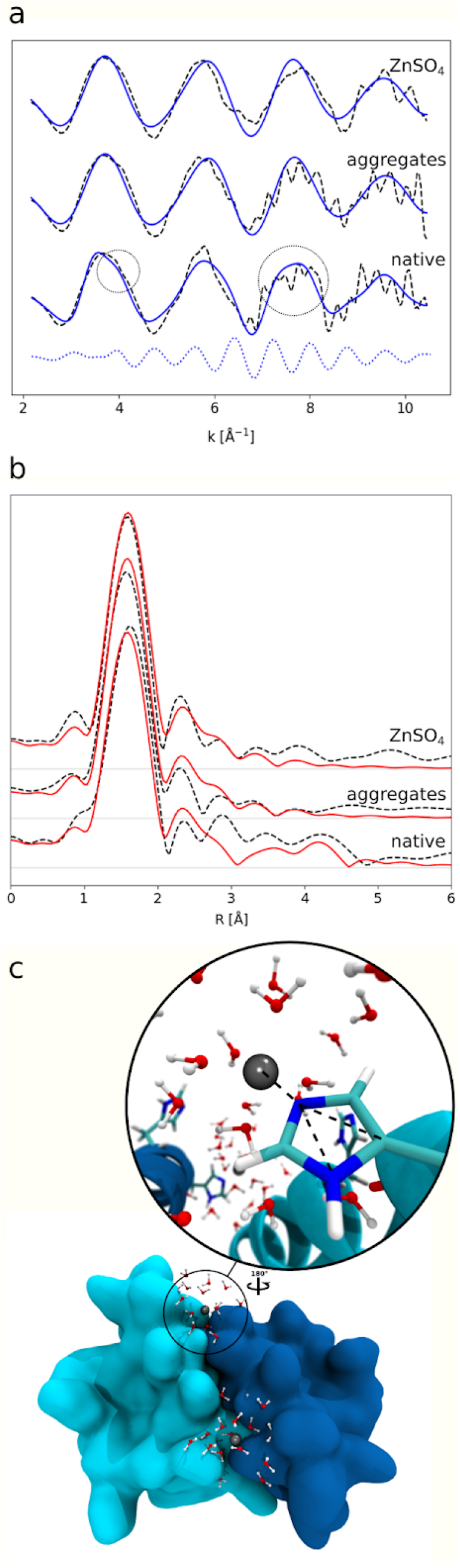
X-ray absorption spectroscopy sheds light on the zinc environment in native insulin and insulin aggregates in solution. **a** k^3^-weighted EXAFS transmission spectra (dashed lines) of a native insulin solution, an insulin solution treated at 60° C for about 24 h (aggregates), and a ZnSO_4_ solution. Blue continuous lines represent the signals simulated on the basis of the structure of a fully solvated Zn^2+^ ion (for the ZnSO_4_ solution and for the aggregates) and of the structure of a solvated Zn^2+^ ion in the presence of a histidine residue (native insulin, see panel c). Details on the structural models are given in the Methods. Blue dotted line represents the component of the simulated signal that is responsible for the features present only in native insulin and indicated by the circles (i.e. a shoulder at 4 Å^-1^ and a broadening of the peak around 7.6 Å^-1^). **b** k^3^-weighted Fourier-transformed EXAFS transmission spectra (dashed lines) of the same samples as in the top panel (**a**). Red continuous lines represent the signals simulated with the same structural models as in the top panel (**a**). **c** The structure of the dimeric human insulin (PDB 5CNY) is represented with a zoom on the structure of the zinc environment used to analyze the EXAFS data. Zinc is represented by a grey bead surrounded by water molecules and an histidine residue. Dashed black lines indicate the multiple scattering contribution from the histidine to the EXAFS signal.

### Dynamic Light Scattering (DLS)

DLS measurements were performed using a DynaPro Nanostar (Wyatt Technology) detector in Wyatt cuvettes with 10 μL of protein solution. Each measurement consisted of 10 times five seconds of integration with auto-attenuation enabled. Insulin was dissolved in 0.25 M NaCl, pH 1.8, H_2_O solution with or without EDTA, then centrifuged at 16000 rpm for 30 minutes. Samples were measured three times and data were analyzed using DYNAMICS software (Wyatt Technology).

### Circular dichroïsm (CD)

Insulin was dissolved in pure H_2_O by lowering the pH with sulfuric acid to avoid chlorine for CD measurement. Stock insulin solution at 2 mg/mL was filtered, then diluted to reach the desired concentration with or without the appropriate amount of EDTA. Measurements were performed on a JASCO J-810 spectropolarimeter at 20 nm/s scanning speed with six accumulations.

### X-ray powder diffraction

A fraction of the insulin-sample powders was used for X-ray diffraction measurements. A small amount was sealed in a Mylar capillary and diffraction patterns were recorded on beamline ID30B at the European Synchrotron Radiation Facility (Grenoble, France) using the following parameters: 20 images with oscillation step of 1° and 0.5 s exposure time for each image (100 % transmission for 4×10^12^ ph/s at λ = 0.98 Å).

### Aggregation kinetics

Insulin was dissolved in 0.25 M NaCl, pH 1.8, H_2_O solution, with or without EDTA, then filtered. 100 μL were pipetted into wells of a 96-well plate and 2 μL of 1 mM Thioflavin T (ThT) was added to each. Each well was sealed using vacuum grease and a glass slide to prevent any evaporation. One measurement was then performed every 5 min at 60°C for 24 h using 450 nm excitation light and a 490 nm detection filter for ThT. The measurements were performed on a BioTek Synergy H4 plate-reader using the bottom configuration and a manual gain of 75.

### Electron microscopy

Samples were absorbed on the clean side of a carbon film, that was then deposited on mica, stained with 2% (w/v) uranyl acetate (pH 4.5) and transferred to a 400-mesh copper grid. The images were acquired under low dose conditions (<10 e^-^/Å^2^) with defocus values between 1.2 and 2.5 μm on a Tecnai 12 LaB6 electron microscope at 120 kV accelerating voltage using a Gatan Orius 1000 CCD camera.

### Optical microscopy

For each sample observed, Thioflavin T was added to a final concentration of 20 μM. Subsequently, 8 μL of the sample were deposited between a glass slide and a cover slip. The samples were observed by an Olympus IX81 microscope equipped with GFP filter cube set and differential interference contrast (DIC) and a sCMOS Hamamatsu Orca Flash4 camera. The control of the microscope and the image acquisition were performed using the software Volocity.

## Results

### Screening and selection of homogeneous aggregated samples

The first step of the present work was to screen and identify the experimental conditions for producing homogeneous samples, i.e. predominantly containing either fibrils or spherulites. This step was necessary to probe any potential difference in protein dynamics depending on the specific aggregate formed. The effect of salt, protein concentration, temperature (which has been shown to affect spherulite size and concentration (14)) and mechanical perturbation on insulin amyloid aggregation was investigated.

#### Effect of salt and protein concentration

Insulin was incubated for 24 h at 60° C in different solutions with salt and protein concentration ranging from 0.1 to 0.5 M and from 0.5 mg/mL up to 5 mg/mL, respectively. Whereas protein concentration has no relevant effect on the final aggregate formation (i.e. both spherulites and fibrils were found at all the concentrations explored), salt concentration strongly influences it: at high salt concentration (□ 0.5 mM), neither spherulites nor fibrils were observed (Fig. S1 in the Supporting Material).

#### Effect of temperature

Insulin was then incubated for 24 h at either 50, 60 or 70° C and at salt and protein concentrations of 0.25 M and 2 mg/ml, respectively. Amyloid aggregation in quiescent conditions (i.e. without shaking) occurs only at or above 60° C (Fig. S2 in the Supporting Material).

#### Effect of mechanical perturbation

Having defined the conditions for maximizing amyloid aggregation in quiescent conditions, mechanical perturbation was used to select either fibrils or spherulites. It was found that shaking the sample at 600 or 900 rpm during incubation produces a sample in which a large majority of the initial native protein is converted into elongated amyloid-like fibrils, even at 50° C. However, even though shaking allows the suppression of spherulites formation, it has a noteworthy effect on the morphology of fibrils. Electron microscopy revealed the presence of well-defined fibrils at 600 rpm and strongly aggregated ThT-positive material at 900 rpm (Fig. S3 in the Supporting Material). A pure spherulite sample could be obtained by heating the protein in 0.25 M NaCl solution at 60° C or 70° C without shaking, and then including a post-production treatment based on a series of vortexing/centrifugations/supernatant removal steps (see Methods for details). After this treatment, no fibrils could be found in the samples (Fig. S4 in the Supporting Material).

The screening described above enabled us to define conditions for obtaining homogeneous fibril and spherulite samples. Fibrils were produced by incubating at 50° C for 24 h a human insulin solution at 5 mg/ml, pH 1.8, 0.25 mM NaCl under shaking at 600 rpm. Spherulites were produced by incubating at 60° C for 24 h a human insulin solution at 5 mg/ml, pH 1.8, 0.25 mM NaCl in quiescent conditions and then isolated by the post-production treatment mentioned above and described in Methods. Both fibril and spherulite samples were then washed in pure D_2_O (see Methods for details) and subsequently freeze-dried. As expected, the macroscopic morphology of both fibrils and spherulites appears slightly damaged after freeze-drying, yet the main features of both fibrils and spherulites remain clearly visible (Figure 1). Moreover, the presence of cross-β structure is revealed in both amyloid samples by X-ray powder diffraction, showing marked differences if compared with the alpha-helix rich native state (Fig. S5 in the Supporting Material).

### Protein dynamical transition is suppressed in insulin fibrils and spherulites

Incoherent neutron scattering of native insulin and of insulin fibrils and spherulites was then employed to measure the elastic incoherent signal as a function of temperature and the momentum transfer *q*. The dependence of the elastic incoherent signal on *q* provides information on the average motions of individual atoms in the system. From the *q*-dependence of incoherent elastic scattering from a D_2_O-hydrated protein powder, one can obtain the ensemble-averaged apparent mean square displacements (MSD) of protein hydrogen atoms, reflecting sub-nanosecond signature of vibrations as well as backbone and side chain relaxation (56). The scattering signal from hydrated insulin powders (hydration level h = 0.4 [D_2_O mass]/[protein mass]) was acquired during a heating scan at 0.2 K/min from 10 to 300 K, yielding MSDs as a function of temperature (Figure 2). Native insulin shows the typical temperature dependent MSDs of hydrated proteins (53), with harmonic behaviour at cryogenic temperature, a first anharmonic onset at ~100 K, attributed to methyl group rotations entering the experimental window (78–80), and a second onset of water-sustained (81, 82) anharmonic motions above ~240 K (the so-called protein dynamical transition (53)). In both amyloid insulin species, i.e. fibrils and spherulites, the protein dynamical transition is completely suppressed, as in a dry protein (78), proving that this property is independent of the type of amyloid aggregates. Interestingly, the MSDs of native insulin treated with EDTA, known to be a highly efficient zinc-chelating agent, display a similar temperature dependence to that of aggregated samples and also lack the protein dynamical transition (Figure 2). Intriguingly, the dynamical transition can be partially suppressed by successive washing steps in pure D_2_O (Fig. S6 in the Supporting Material). As the formation of crystalline ice within the sample might also lead to dewetting of the protein at sub-zero temperatures, and hence to the observed dry-like MSD temperature dependence, we measured the neutron diffraction signal of insulin fibrils and spherulites to determine if crystalline ice had formed at low temperature. As shown in Fig. S7 in the Supporting Material, no Bragg peaks of hexagonal ice are present in the diffraction pattern of fibers at 200 K. A broad peak centered at ~1.65 Å^-1^ and a peak at ~1.35 Å^-1^ indicate the presence of amorphous water and of amyloid cross-β structure, respectively.

### Zinc release affects the kinetics of insulin amyloid aggregation

As revealed by neutron scattering, insulin amyloid species (both fibrils and spherulites) and EDTA-treated insulin (where zinc is expected to be chelated by EDTA and no amyloid aggregation is induced as shown by X-ray diffraction data in Fig. S5) are characterized by the same dynamical behavior. This suggests that the presence/absence of zinc may affect the aggregation kinetics. To verify this, we monitored insulin aggregation kinetics at 60° C as a function of EDTA concentration. Thioflavin T (ThT, a probe whose fluorescence correlates with amyloid formation (83)) fluorescence was used as a probe of aggregate formation. As shown in Figure 3, the kinetics in the absence of EDTA and without mechanical shaking is characterized by a double sigmoidal curve as previously observed (84): a lag phase is followed by a first increase of the fluorescence signal, then by a plateau prior to a second increase. Adding EDTA clearly reduces the double-sigmoidal trend, which is essentially suppressed at high EDTA concentration. As expected, optical and electron microscopy observations at different time points along the aggregation kinetics showed that fibrils and spherulites are formed roughly at the same time, with more numerous and larger spherulites appearing in the presence of EDTA (Figure 3). As shown above, shaking the insulin solution during incubation prevents spherulite formation. The kinetics experiments were then repeated as for those shown in Figure 3 but in the presence of mechanical shaking, which inhibits the formation of spherulites, and results in pure fibril samples. The results are shown in Fig. S8 in the Supporting Material and reveal that shaking suppresses the first plateau phase, while the addition of EDTA in the presence of mechanical perturbation does not produce any relevant effect. In addition, no spherulites were found with or without EDTA with mechanical shaking. In order to explore any possible structural/conformational perturbation directly induced by EDTA, we also verified that in the range of EDTA concentration used both the oligomeric state and secondary structure of native insulin at room temperature are largely unaffected. In fact, a dynamic light scattering (DLS) analysis of size distribution showed a monodisperse signal for native insulin, with a single peak at a radius of approximately 2.8 nm (suggestive of the presence of an insulin dimer), barely affected upon addition of EDTA up to a concentration of 5 mM (Fig. S9 in the Supporting Material). Circular dichroism (CD) in the 200-260 nm spectral region indicated that insulin secondary structure content is unchanged upon EDTA addition even at high protein:EDTA ratios (Fig. S10 in the Supporting Material).

### Insulin releases zinc upon aggregation

In the light of the effect of EDTA on both insulin dynamics and aggregation kinetics, our hypothesis is then that, upon aggregation, insulin molecules release the zinc ions. To test this hypothesis, the presence of zinc in the different samples was explored. μPIXE (59, 60) analysis was first used to determine the zinc concentration in the insulin samples investigated by neutron scattering. The results are shown in Figure 4 and stoichiometric ratios are summarized in Tab. S1 in the Supporting Material. μPIXE analysis revealed that native insulin contained 1.5 zinc atoms per insulin monomer, while zinc was below the detection limit (<0.001 atoms of zinc/monomer) in both fibrils and spherulites and, as expected, most of the zinc was removed in the EDTA-treated sample (0.02 zinc atoms per insulin monomer). However, the washed insulin sample still contains zinc with a stoichiometry of 0.116 zinc atoms per insulin monomer.

To get insights into the zinc-insulin interaction in native and amyloid conditions in solution we used XAS spectroscopy. XAS can be divided into X-ray absorption near-edge spectroscopy (XANES) and extended X-ray absorption fine structure (EXAFS), with XANES providing information on the oxidation state, geometry and electronic configuration of the absorber atom. In contrast, EXAFS allows solving an atom’s local environment in terms of the number of neighboring atoms, their distance from the absorber atom and their disorder on a short length scale. XAS experiments were performed at the zinc K-edge (9660.75 eV) on both a native insulin solution and an insulin solution incubated at 60° C to induce the formation of amyloid species (since in quiescent conditions both fibrils and spherulites are always formed, as described above, this sample was named “aggregates”), and on a ZnSO_4_ water solution. The XANES spectra in Figure S11 in the Supporting Material do not show any significant difference between the various samples, indicating that Zn^2+^ ions are solvated in all the samples. Figure 5 (panel a) shows k^3^-weighted EXAFS transmission spectra and their k^3^-weighted Fourier transforms (panel b) for the three samples. The experimental curves in real space (Figure 5b) are comparable up to about 3 Å, indicating that the first shell is very similar in all three samples. At larger distances, the curve for native insulin shows small but significant differences in the 3-5 Å range compared with the other two samples (in particular the peak at ~4.2 Å present only in native insulin). Small but significant differences are also evident in the reciprocal space (Figure 5a), namely a shoulder at 4 Å^-1^ and a broadening of the peak around 7.6 Å^-1^ visible only in native insulin. Figure 5a shows fitted curves in terms of structural models (see Methods). The structural models contain either only a solvated Zn^2+^ ion, used to analyze data of both the ZnSO_4_ and the amyloid insulin solution samples, or a solvated Zn^2+^ ion in the vicinity of a histidine residue (Figure 5c), known to coordinate zinc in insulin, used to analyze data of the native insulin solution. Both models were obtained from constrained molecular dynamics simulations, based on refined geometrical parameters (85) and used to extract coordinates that were processed by GNXAS, as described in the Methods. Structural parameters obtained from this analysis are reported in Tab. S2 in the Supporting Material. A good agreement was found between experimental and calculated curves (Figure 5a). The Fourier-transformed calculated curves also reproduced the peak positions of the Fourier-transformed experimental data up to ~4.5 Å, in particular the differences in the 3-5 Å range (Figure 5b). XAS results indicate that Zn^2+^ is coordinated with a histidine residue in the native insulin solution while it is released in solution upon amyloid formation.

## Discussion

Our study aimed at comparing the molecular dynamics of native and amyloid insulin species by neutron scattering and complementary biophysical methods. Zinc binding proved to be pivotal in controlling not only molecular dynamics at mesoscopic scale but also the aggregation kinetics. To enable the comparison of various insulin species and monitor potential differences in the protein dynamics depending on the type of aggregates, we first established protocols to produce batches containing either native insulin, insulin fibrils or insulin spherulites at a high level of homogeneity. We found that while fine-tuning temperature and salt concentration allows amyloid aggregation into both fibrils and spherulites to be observed, shaking the vial during incubation leads to fibril formation and full inhibition of spherulite formation. We can argue that this inhibition of spherulite formation originates from mechanical perturbation preventing the formation of sufficiently stable spherulite-specific precursors. Spherulites are always produced along with fibrils in quiescent conditions. However, taking advantage of the much larger sedimentation rate of spherulites compared to the other species, they could be isolated from fibrils using gentle centrifugation. Neutron backscattering spectroscopy showed a drastic change in the dynamics between native insulin and insulin in aggregated form above ~240 K, but no significant difference between the two types of aggregates (spherulites and fibers) were observed (Figure 2). Indeed large-amplitude motions associated with the protein dynamical transition are fully suppressed in insulin amyloid fibers and spherulites, so that the temperature-dependence of the mean square displacements resembles that of completely dry, non-functional proteins. In the native-insulin control measurement, a dynamical transition takes place (Figure 2). These results are in contrast with our previous results on tau protein (52), where no differences between native-state and fibril dynamics were observed on the same timescale, and on concanavalin A, where internal dynamics were found to be larger in the amyloid than in the native species (51). The tau results were suggestive of the protein internal energy depending only on the amino acid composition (86) and not on the structure or aggregation state, as also supported by solution studies of protein denaturation showing that the internal dynamics remain unchanged upon denaturation (87).

In order to rationalize the differential dynamical behavior of native and amyloid insulin, we suspected that interaction with Zn^2+^ ions might play a role, since zinc is known to be essential for insulin expression and activity (33). We investigated how zinc binding affects the kinetics of insulin amyloid aggregation. Upon thermal treatment of the insulin solutions, ThT fluorescence profiles first showed an increase in the signal barely dependent on zinc presence, which has been suggested to be related to the formation of prefibrillar aggregates or early formation of fibrils (84) small enough to leave turbidity and temperature dependence unaffected. We attributed the intermediate plateau phase to spherulite formation as shown by fluorescence and electron microscopies (Figure 3): both fibrils and spherulites are present, with spherulites being larger and more numerous in the presence of EDTA, i.e. when zinc is removed. After the first plateau a second increase occurs, followed by a final plateau, where spherulites are densely present and significantly larger. Increasing EDTA concentration gradually suppresses the double-sigmoidal behavior and enhances spherulite formation. This indicates that the presence of zinc is the limiting step for the more efficient conversion of insulin native molecules into spherulites. When the zinc is fully removed by adding a sufficient amount of EDTA, the intermediate phase disappears. If the loss of zinc occurs at the level of single molecules prior to aggregation or happens once the molecules are embedded in high MW species remains to be clarified. Importantly, shaking also suppresses the double-sigmoidal behavior as well as the spherulite formation. From the kinetic results we can conclude that: 1) the first plateau can be attributed to spherulite formation; 2) zinc binding inhibits spherulite formation, probably through electrostatic repulsion.

To obtain quantitative information on zinc content in our samples, we performed measurements with the μPIXE technique, the results of which unambiguously show that zinc was below the limit of detection (<0.001 zinc atom/monomer, Tab. S1) in amyloid aggregates, where the dynamical transition is suppressed. Furthermore, an EDTA-treated control sample is shown by μPIXE to have a zinc content reduced by a factor of 75 compared with the native protein (Figure 4), and lacks the protein dynamical transition at 240 K in neutron scattering experiments (Figure 2). We note that the EDTA-treated sample and the native form present the same secondary structure according to CD data (Fig. S10) and are predominantly dimeric according to DLS (Fig. S9). This means that both the conformational and colloidal stability of the solution are not significantly influenced by the presence of EDTA and the loss of zinc is mainly affecting the dynamics of the single protein molecule. Indeed, an insulin sample with a zinc content intermediate between those of native and amyloid insulin (as confirmed by μPIXE, Tab. S1), obtained by successive washing steps of a native sample, shows a dynamical transition but with reduced MSD if compared to the native sample (Fig. S6). Zinc binding to insulin thus modulates molecular dynamics and its absence suppresses functionally relevant motions activated at the dynamical transition in amyloid insulin, without modifying secondary, tertiary or quaternary structure. XAS data in solution (Figure 5) confirmed that zinc is coordinated with histidine residues in native insulin, while it is released upon amyloid aggregation. The XAS results thus established that zinc loss is not an effect of the sample treatment (i.e. buffer exchange and freeze-drying) preceding neutron scattering experiments.

Translating our *in vitro* results to a physiological context, they imply that amyloid aggregation of insulin *in vivo* could induce an accumulation of free zinc ions at the injection sites, where IDA is observed. Zinc is anyway released from monomeric insulin during its function (33), yet our finding suggests that a substantially larger quantity of zinc may be released and could accompany IDA. Zinc overload is known to be a pro-oxidant condition leading to oxidative damage of biomolecules. When free zinc ion concentrations increase, zinc can bind to proteins that otherwise would not interact with zinc and affect their functions, thus inducing the production of free radicals by inhibiting antioxidant enzymes and the mitochondrial respiratory chain (88). On the basis of our results we are not able to quantitatively assess the potential cytotoxicity of zinc release during IDA, because the threshold at which zinc ion concentration becomes high enough to produce pro-oxidant/cytotoxic effects depends on the zinc buffering capacity of cells (33, 88). Future *in vivo* investigations could clarify the physiological impact of the zinc accumulation revealed by our study.

An important information provided by the XAS results is that even in native insulin, which coordinates zinc, the metal ion environment is largely composed of water molecules. The presence of the high charge on the Zn^2+^ ion is known to perturb water molecules in the first and second solvation shells, which correspond to at least 20 water molecules interacting with one Zn^2+^ ion (89). Considering the 1.5 Zn/protein ratio indicated by the μPIXE analysis, the presence of zinc on the protein surface corresponds to a minimum hydration level h = [20 x 1.5 x water MW]/[insulin MW] ≃ 0.1. In other words, the presence of Zn^2+^ favors the hydration of the highly hydrophobic insulin surface, thus influencing the dynamical behaviour as revealed by neutron scattering.

Taken together, the results from neutron scattering, μPIXE, and XAS clearly show that zinc binding is the key parameter which allows hydrated insulin to undergo the protein dynamical transition, while the dramatic dynamical effect on fibrils and spherulites has to be attributed to loss of zinc. Removal of zinc results in a complete loss of water-sustained backbone and side-chain motions with the MSD profile corresponding to that of a dry protein, thus suggesting a dehydration effect upon zinc release. As previously suggested (52, 90, 91), the changes in hydration water entropy may be one of the ‘driving forces’ leading proteins to form amyloid aggregates. Restructuring of hydration water is expected to be involved both in the initial steps of unfolding/crossβ-amyloid formation and in the growth of amyloid aggregates, thereby implying a possible change in the water configurational entropy (90). Protein dehydration has also been shown to modify amyloid aggregation kinetics (92). The presence of metal ions integrated within the protein structure, which generally influences protein solvation and electrostatics of the protein surface (93), can add a key parameter to the energetic balance between native and amyloid state. In the case of human insulin, which is highly hydrophobic(94), we postulate that zinc binding favors protein hydration, thus modifying both protein dynamics, as revealed by neutron scattering, and water entropy. Indeed, dewetting of hydrophobic groups (which are known to reduce water mobility (95) and entropy (96)) during aggregation, and subsequent release of water molecules in the bulk phase, corresponds to an increase of water entropy (91). The presence of zinc coordinated to the protein reduces the hydrophobicity of the insulin surface and then the propensity for aggregation, whereas zinc removal increases the entropy difference between native and amyloid state and then promotes amyloid aggregation.

It has been shown that a Thr to His mutation in the chain A of human insulin leads to a significant increase in the lag phase of amyloid aggregation (97). More generally, studies of protein rational design (48, 98) have shown how the introduction of His binding sites for Zn^2+^ into hydrophobic peptides can modify both their stability and dynamics. Our results suggest that protein hydration changes upon Zn^2+^ binding/release can contribute to such modifications, as revealed by our finding that both dynamics and aggregation kinetics of human insulin change as an effect of Zn^2+^ release from His binding sites. Changes in metal-modulated protein hydration might be a general strategy to control protein stability and tune protein aggregation into amorphous and ordered super-structures.

## Author contributions

K.P., V.F., M.W., and G.S. designed the research; K.P., G.S., D.N., and M.Z. performed neutron backscattering spectroscopy and K.P. analyzed data; K.P. and V.C. performed neutron diffraction and analyzed data; K.P. and V.F. defined sample preparation conditions and K.P. prepared and characterized the samples; G. W. G. and E. F. G. performed μPIXE and analyzed data; A.L., K.P. and G.S. performed XAS and A.L. analyzed data based on structures extracted from MD simulations performed by K.P.; K.P. performed CD, DLS and fluorescence spectroscopy; K.P. and G.S. performed X-ray powder diffraction; K.P., V.F., and G.S. wrote the paper with input from all authors.

The authors declare no competing interests.

## Acknowledgments

This work used the platforms of the Grenoble Instruct-ERIC Center (ISBG: UMS 3518 CNRS-CEA-UGA-EMBL) with support from FRISBI (ANR-10-INSB-05-02) and GRAL (ANR-10-LABX-49-01) within the Grenoble Partnership for Structural Biology (PSB). V.F. acknowledges VILLUM FONDEN for the Villum Young Investigator Grant “*Protein Superstructures as Smart Biomaterials (ProSmart)*” 2018-2023 (project number: 19175).

The electron microscope facility is supported by the Rhône-Alpes Region, the Fondation Recherche Medicale (FRM), the fonds FEDER, the Centre National de la Recherche Scientifique (CNRS), the CEA, the University of Grenoble, EMBL, and the GIS-Infrastructures en Biologie Santé et Agronomie (IBISA).

The financial support provided by JCNS to perform neutron scattering measurements at the Heinz Maier-Leibnitz Zentrum (MLZ), Garching, Germany is gratefully acknowledged. The access to ILL and ESRF beamlines to perform diffraction characterization is also acknowledged.

We thank Daphna Fenel, Christine Moriscot and Guy Schoehn from the Electron Microscopy platform, Caroline Mas and Marc Jamin for assistance and access to the Biophysical platform, Aline Le Roy, Michel Thépaut and Christine Ebel for assistance and access to the Protein Analysis On Line (PAOL) platform.

